# Developing a Sensory Representation of an Artificial Body Part

**DOI:** 10.1101/2025.06.16.658246

**Authors:** Lucy Dowdall, Maria Molina-Sanchez, Giulia Dominijanni, Edmund da Silva, Viktorija Pavalkyte, Ema Jugovic, Matteo Bianchi, Fumiya Iida, Dani Clode, Tamar R. Makin

## Abstract

Somatosensory feedback is essential for motor control, yet artificial limbs are thought to lack such feedback. We investigated how the body and brain gather informative sensory signals from a wearable augmentation interface (a robotic digit for motor augmentation), and whether naturally-mediated feedback can support technological embodiment.

Participants intuitively interpreted natural feedback across perceptual tasks, performing comparably to state-of-the-art artificial feedback systems. fMRI revealed an immediate and distinct, topographically-organised representation of the robotic digit. After longitudinal training, this representation was further refined, becoming more similar to the biological digits, which correlated with increased subjective somatosensory embodiment.

Our findings demonstrate that wearable devices naturally provide a powerful source of feedback which is immediately integrated with our body. Long-term use can then promote device-embodiment across the sensorimotor hierarchy.

## Introduction

Motor control is thought to be governed by the sensorimotor loop (*1*), where sensory feedback informs about the success of our actions to drive motor learning. Somatosensory feedback (primarily encompassing touch and proprioception) plays a crucial role in both online motor planning and error correction, as well as offline skill acquisition and consolidation (*2–4*). Its importance is strikingly demonstrated in the motor deficits observed in its absence (*5*,*6*).

This reliance on somatosensory feedback poses a challenge for technologies that compliment, replace or expand human movement abilities, such as prosthetic limbs (*7*,*8*), brain-machine interfaces (*9*,*10*), and augmentative extra robotic body parts (*11–15*). Such technologies are said to suffer from ‘open-loop’ control (*16–18*), where motor signals are sent out, but no corresponding somatosensory feedback is received. To address this, research has focused on artificial sensory feedback systems, aiming to replace this missing input.

Primarily focusing on touch, current efforts to produce artificial feedback include exploration of invasive electrical stimulation at the periphery (*19*,*20*), spinal cord (*21*,*22*) and primary somatosensory cortex (S1) (*23*,*24*), in addition to non-invasive electrotactile (*25*,*26*), mechanotactile (*27–30*) and vibratory feedback signals (*31–33*) (reviewed in *34*). However, unlike the rich, multimodal information conveyed in natural touch, these solutions offer signals with reduced dimensionality that cannot replicate the same diversity of feature extraction, raising questions regarding the fidelity of artificial haptics (*35*). Moreover, artificial signals are designed to meet perceptual thresholds, thus risking overriding more subtle yet highly informative existing sensory inputs. This highlights a particular barrier for technologies that extend, rather than replace, motor abilities. We have limited neural capacity that is already fully dedicated to representing our biological body (*36–38*). Consequently, these devices must share existing resources, without disrupting natural motor control (soft embodiment) (*39*,*40*).

Therefore, a key guiding principle is how artificial systems feedback may be best ‘embodied’ into our sensorimotor system. Embodiment is an umbrella term that broadly captures the ability to relate to external objects as if they were a body part (*41*). Theoretically, achieving embodiment requires a closed-loop system, making sensory feedback essential (*42*). However, how integration into the sensorimotor system manifests at the neural level remains unclear (*43*,*44*).

We aimed to determine how the sensory representation of an extra robotic body part first emerges and is then refined through sensorimotor experience. We used the Third Thumb (Dani Clode Design; Figure 1A; hereafter Digit 6 (D6)), a supernumerary robotic digit designed for collaborative use with the biological fingers to extend hand functionality. We have previously demonstrated that people can successfully perform a proprioceptive task with D6, despite not having any additional artificial feedback (non-sensorised) (*45*,*46*). Similar results were also replicated with non-sensorised prosthetic devices (*47*, see also *48*). This suggests that artificial limbs may not be inherently open-loop systems, and naturally-mediated somatosensory cues from how devices directly interact with our somatosensory system may be leveraged to provide somatosensory awareness and help close the sensorimotor loop. Specifically, inputs relating to D6’s movement and interactions travel via the physical connection of D6 with the palm (Figure 1B). Similarly, naturally-mediated somatosensory feedback from the controllers of D6 have been shown to support motor learning (*49*).

**Figure 1:**
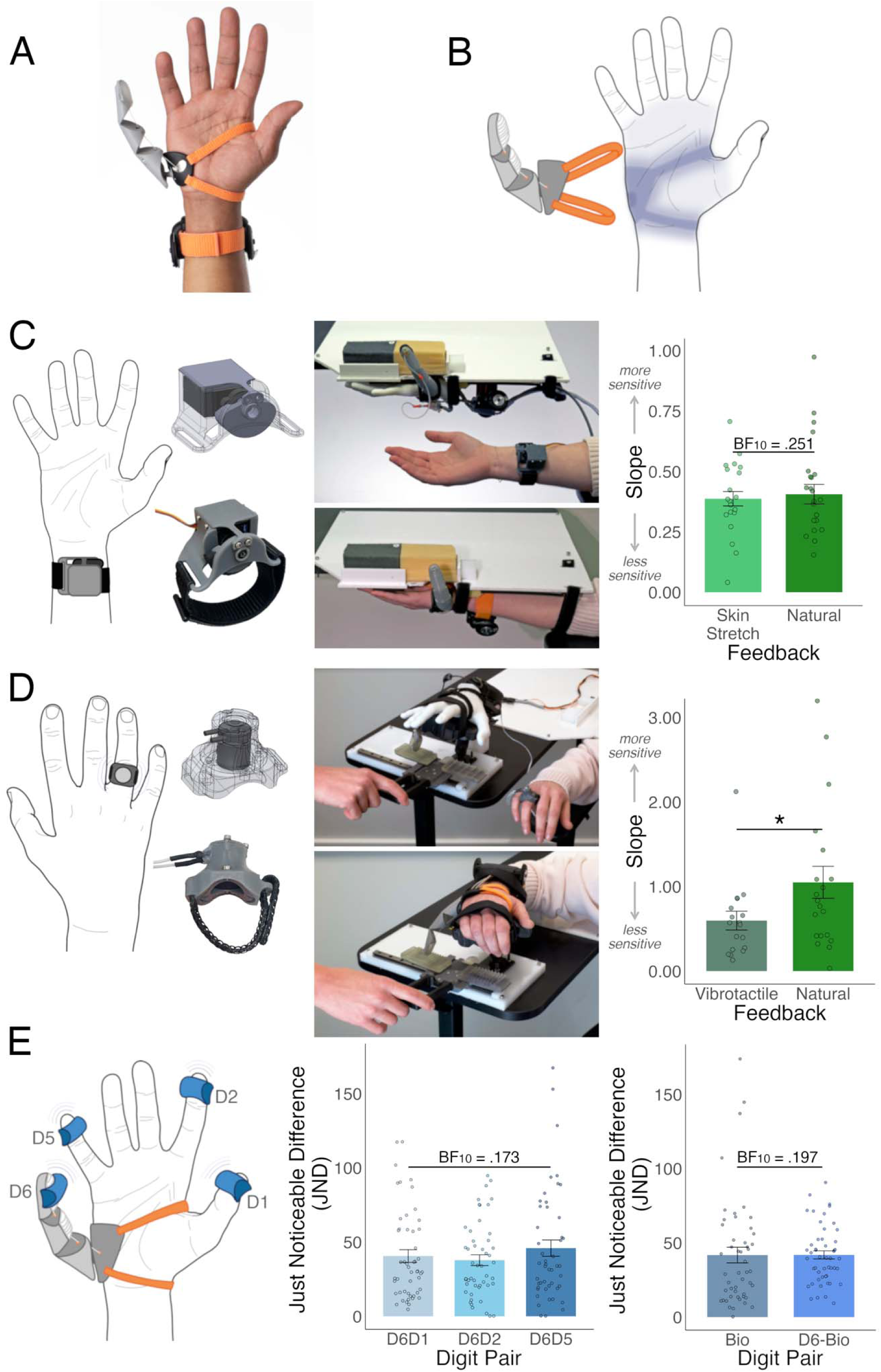
Users can easily extract detailed sensory information to inform perceptual judgements with an artificial body part. (A) The Third Thumb (Dani Clode Design; Digit 6 (D6)) is a 3D-printed extra robotic digit, worn on the ulnar side of the hand and controlled by force sensors fixed underneath the big toes. (B) Sensory feedback is passively received on the side of the hand and distributed across the hand from how D6 is worn and moves on the hand, even without the active mediation of sensors and actuators; hereafter “natural” feedback. (C) Incorporation of Skin Stretch artificial feedback into the D6 design, where deformation of a pressure sensor in the D6 tip produced a proportional amount of linear skin stretch in the inner wrist (left). Participants performed just as well in a material deformation task (middle) with the Skin Stretch feedback as with the Natural feedback (BF_10_ =.251, right). (D) Incorporation of Vibrotactile artificial feedback into the D6 design, where non-intentional displacement of the D6 tip produced a proportional amount of Vibrotactile feedback delivered to the ring finger (left). The Natural Feedback outperformed the Vibrotactile Feedback (right) in a texture discrimination task (middle). (E) In a temporal order judgement (TOJ) task, motors were attached to the biological thumb (D1), index finger (D2), little finger (D5) and (non-sensorised) D6 and were used to stimulate digit-pairs in quick succession (left). Participants could successfully make spatiotemporal judgements about the D6 digit-pairs, performing similarly to when performing the task with only their biological fingers (right). Participants also performed similarly across all D6 digit-pairs, regardless of distance between D6 and the biological digit (middle). * denotes p < .05.

In the current study, we first characterised how “natural” sensory feedback on the hand can support sensory perception of a robotic extra digit in comparison to state-of-the-art artificial sensory feedback. We will refer to this feedback as ‘natural’, even though the source of the input is artificial (e.g. external stimulation delivered to D6 through its interactions with the environment), as the input is mediated from D6 to the hand (and nervous system) without the support of sensors and actuators. We then examined how this natural feedback integrates with the somatosensory representation of the hand at both the perceptual and neural levels. Finally, we explored construction of a sensory representation of D6 in relation to the human hand following extended motor training. We find that the brain rapidly accesses and integrates natural tactile feedback into sensory representations across the sensorimotor hierarchy, offering a potential means to ‘close’ the sensorimotor loop for wearable technology.

## Results

### Natural feedback signals can be easily and intuitively interpreted by a novice user

To demonstrate the potential utility of natural feedback, we first developed a proof-of-concept test to index natural somatosensory perceptual abilities. We isolated two components that contribute to the tactile and proprioceptive somatosensory awareness of the Third Thumb (Digit 6; D6): skin stretch from D6 movement and vibrotactile input from D6-object contact. Two artificial feedback systems were developed to replicate these aspects, enabling a perceptual comparison of natural and artificial touch.

To deliver artificial skin stretch, we developed a feedback system, where deformation received on the tip of D6 produced a proportional amount of linear skin stretch via an actuator worn on the user’s inner wrist (Figure 1C). For the vibrotactile feedback system, any displacement of D6, due to object impact, produced a proportional amount of vibration delivered via an actuator (*50*) worn on the user’s ring finger (Figure 1D). We tested each of these systems on a material discrimination task: material deformation for the skin stretch system (Figure 1C), and texture coarseness for the vibrotactile system (Figure 1D). In a within-participant design (*N*=22), performance was compared relative to the wearable D6 device which was originally designed to mediate these sensory components via natural touch (i.e. sensory information that is naturally mediated to the hand without the aid of sensors and actuators; Figure 1B). In each trial, feedback resulting from interactions with two different materials was presented to the (blindfolded) participant (either remotely via the artificial system, or directly via the worn D6). Individual participants’ responses (two-alternative force choice; 2AFC) were fitted to a psychometric curve for each condition, and the slope was extracted as a measure of discrimination ability. Larger values (steeper slope) indicated increased sensitivity for discriminating between materials.

We found that, as expected, participants were able to successfully perform each of the tasks using the relevant artificial feedback system (slope significantly above zero, *W*≥171,*p*<.001). Participants were also able to perform the task with just the natural feedback extracted from the worn D6 (*W*≥190,*p*<.001), demonstrating that natural feedback signals can be easily and intuitively interpreted by a novice user. Crucially, participants performed comparably with the natural and artificial systems (skin stretch: *W*(19)=88,*p*=.546,BF_10_=.251), and even outperformed the vibrotactile artificial system (*W*(15)=111,*p*=.025). This set of results demonstrates that natural feedback–when D6 is worn–provides the same level of perceptual detail as the bespoke artificial systems, but also provides the versatility to mediate different sensory components across material demands in an integrated system.

As a control, we also indexed perceptual abilities with D6 against natural sensing abilities, where participants performed a variation of both discrimination tasks with the biological thumb (D1) (*N*=10). Participants consistently outperformed the natural D6 feedback (*U*>154,*p*<.017). Therefore, despite the impressive abilities of the natural feedback, it is still not comparable to natural sensing abilities.

### “Natural” feedback can be integrated with sensory information from the biological hand

Given D6 is designed to be used in collaboration with the biological hand, we next aimed to index how the somatosensory system integrates coordinated somatosensory inputs across D6 and the biological digits.

In a new set of participants (*N*=50; novice D6 users), we employed a Temporal Order Judgement (TOJ) task to determine if the natural sensory feedback received when wearing D6 is integrated in the same spatiotemporal reference frame as the biological fingers. Motors were worn on the tips of paired digits to deliver vibrotactile stimuli with various interstimulus intervals (four digit-pairs: D6 & biological thumb (D1), D6 & index finger (D2), D6 & little finger (D5), and D5&D1). Responses (2AFC) were fitted to a psychometric curve for each digit-pair, where the just noticeable difference was extracted as a measure of discrimination ability (see supplementary material for point of subjective equality as a measure of bias).

We compared D6 digit-pairs performance to the biological-pair. Participants could not only perform the task with D6, but performed just as well as with their biological hand (*W*>506,*p*>.661,BF_10_≤.194; Figure 1E). This successful integration implies the same spatiotemporal reference frame is being used to extract the relevant features from the stimuli. We then examined whether inputs closer to D6 would be more easily confused (as with the biological fingers (*51*,*52*)). However, participants performed just as well across all three D6-pairs (*F*(2,92)=.947,*p*=.392,BF_10_=.173; Figure 1E). This absence of topographical organisation demonstrates the distinctiveness of D6’s sensory information. Together, these results suggest the presence of a distinct and accessible sensory representation for D6.

### Emergence of a neural representation of a robotic finger in novice users

To investigate how a sensory representation of an extra robotic body part may spontaneously emerge, we explored the neural response in the S1 hand area while tactile stimulation was applied to D6 in novice users (*N*=50; Figure 2A).

**Figure 2:**
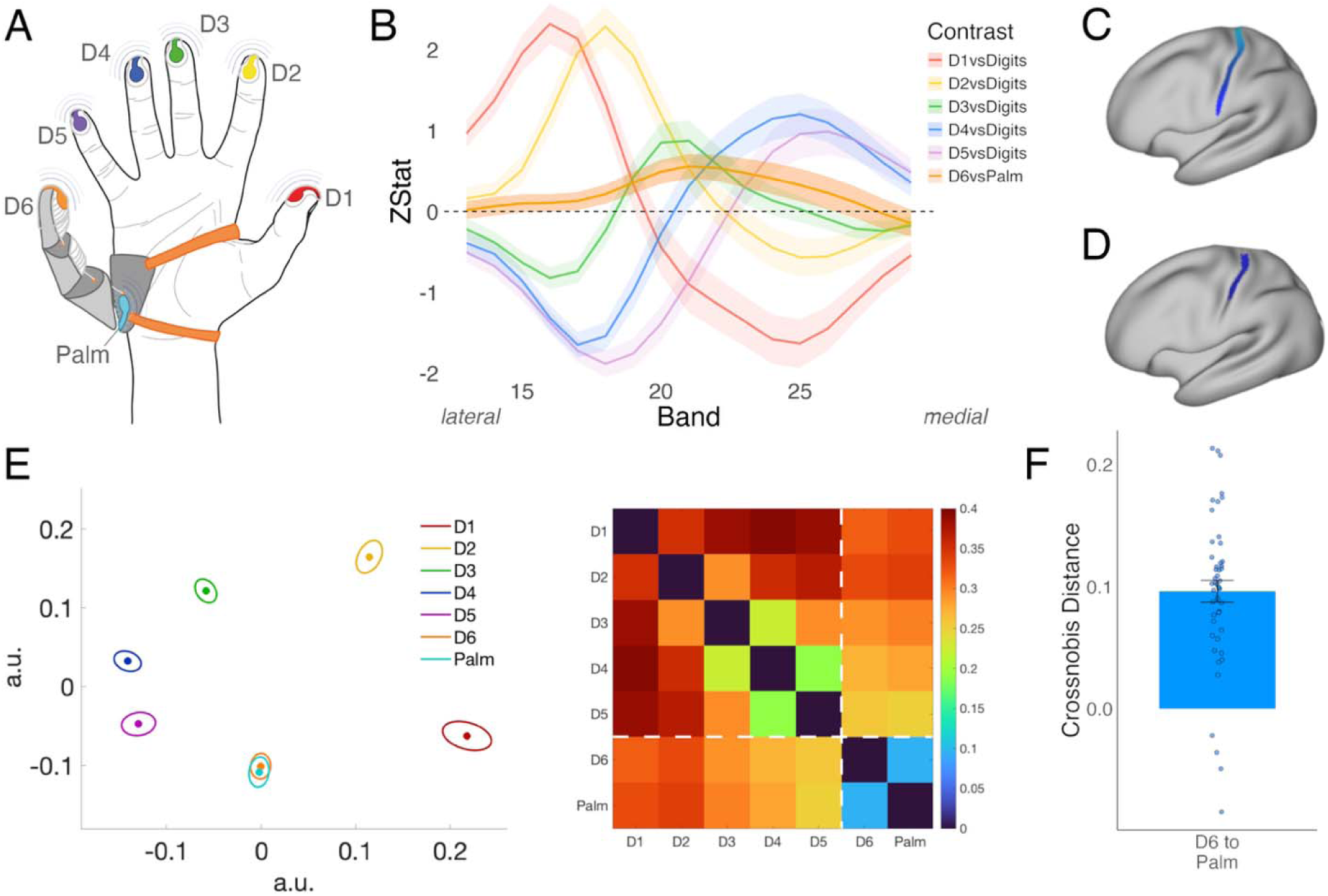
Emergence of a sensory representation of D6 in novice users. (A) Silicone vibrotactors used to deliver stimulations to each of the biological digits, D6 (the Third Thumb), and the side of the hand where the natural feedback is received (the ‘palm’) (left) during the fMRI scan (right). (B) Averaged activity (zstat) elicited along the S1 hand area (lateral (left) to medial (right) for the biological digits and D6 (selectivity; relative to their respective controls - the average of the other biological digits, and the palm of the hand;). (C) S1 (BA3b) hand area region of interest, segmented into 17 equally-spaced bands for line analysis. (D) S1 (BA3b) region of interest for RSA, approximately 2cm proximal/distal to the hand knob. (E) Multidimensional Scaling (MDS) plot (left) and representational dissimilarity matrix (RDM) showing the crossnobis distances between the stimulation conditions, demonstrating the topographical organisation of the hand, and the integration of D6 into this topographical structure. Error bands in MDS represent variability when projecting into 2D space. (F) Crossnobis distance between D6 and the palm is significantly above zero, implying unique information content within the D6 representation even at baseline.

We first visualised the activity elicited in S1 (specifically BA3b) for each of the digits (D1-D6) (Figure 2B). We extracted the average z-values along the hand area (lateral-left to medial-right) and identified the expected somatotopic structure of the five biological fingers (D1–most lateral peak, D5–most medial peak). D6 stimulation also elicited activity within the hand area, peaking near the centre of the hand representation (adjacent to the D3 peak). This indicates that when mediating sensory information from D6, somatosensory cortex receives diffused inputs from across the entire hand.

To examine if a more structured topographic organisation underlies this diffused input, we employed multivariate representational similarity analysis (RSA) to compare D6’s neural representation to that of the palm and biological digits. Our findings reveal that the D6 representation is most similar to that of the palm, reflecting a topographic organisation (see comparable dissimilarity matrix for D6 and Palm in Figure 2E). However, the information content associated with D6 is nevertheless distinct from the palm, demonstrated by a significant cross-nobis distance between the D6 and palm representations (Figure 2F; *W*(48)=1166,*p*<.001). These results provide the first evidence towards a topographic signature of D6, and highlight its unique information content in S1.

### Longitudinal D6 training to promote experience-based plasticity

We next aimed to understand how the somatosensory representation of D6 is refined through sensorimotor experience. We designed a seven-day D6 training regime (Figure 3A), focused on D6-biological hand collaboration (*n*=30). Our control group replicated this week of altered finger usage by training to play the piano keyboard (no D6 involved, *n*=20). The training protocols resulted in extensive improvements in motor performance with D6 during a range of collaboration tasks (main experimental group; as detailed in *46*; Figure 3B), as well as extensive improvements in the control group keyboard task (piano day 1 vs day 7 performance: *F*(1,58)=59.79,*p*<.001; Figure 3C).

**Figure 3:**
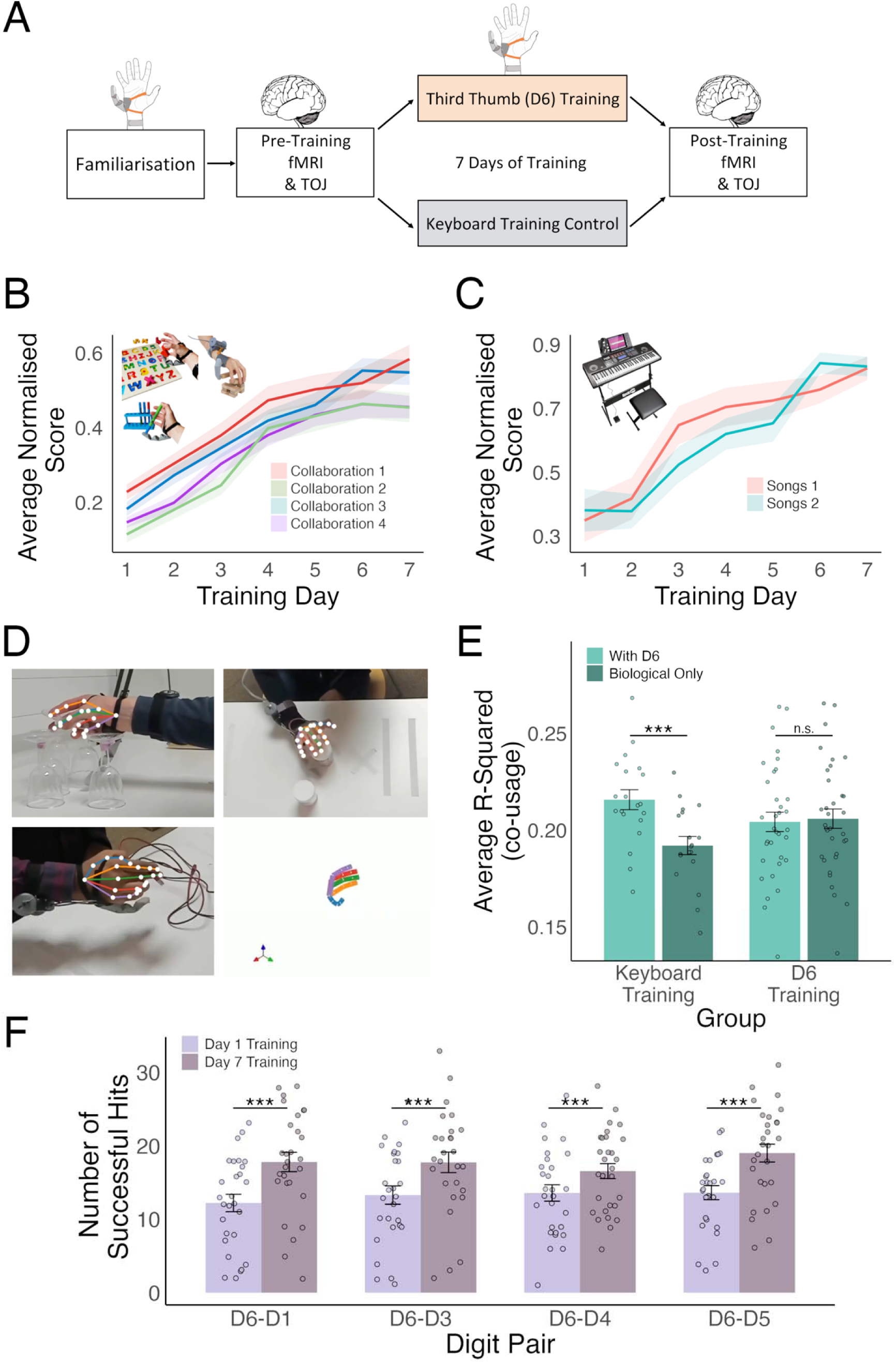
D6 Training. (A) Longitudinal training study design. Participants all underwent a short familiarisation session with D6, followed by the TOJ psychophysics task and passive fMRI scan described above. Participants were then allocated to the D6 Training (experimental group) or Keyboard Training (control group). After their seven days of training, participants returned for another fMRI scan (replicating the first) and performed the TOJ task. (B) D6 training focused on D6-biological hand collaboration-based tasks, exploring different aspects of this motor skill. Participants show large improvements in motor task performance over their week of training. (C) Control group trained to play the piano keyboard over the week of training. Participants showed large improvements in task performance over the week. (D) View from each of the three cameras used to track participants’ hand during the free-choice tasks, with projections of the 2D joint coordinates of the five biological digits, and 3D reconstructed joint angles. (E) When looking at patterns of finger co-usage following training using 3D kinematics (through a markerless tracking setup), Keyboard participants show significantly more individuation when performing ecological tasks without wearing D6 (whilst the D6 training group saw no difference). Averaged over joint and digit-pair for visualisation purposes only. (F) D6 training participants performed a coordination task involving opposing D6 with an individual digit over the course of one minute. Regardless of distance between D6 and the biological digit, similar improvements in coordination were seen across the whole hand. *** denotes p < .001.

Following D6 training, we examined perceptual and motor integration across individual D6-biological digit-pairs. We found that while perceptual integration remained unchanged (likely due to ceiling effects in TOJ performance; pre vs post-training: BF_10_=.23), motor coordination improvements were equally observed across all digits (*F*(1,166.91=50.68,*p*<.001, Figure 3F index finger untested). This indicates distributed sensorimotor gains across the whole hand.

### Refinement of the sensory representation following D6 training

We next examined post-training activity within the S1 hand area. During D6 stimulation, we found the activity peak to be qualitatively higher and more defined, relative to the control group (Figure 4A). This hints at differences in the S1 readout of this peripheral stimulation following training.

**Figure 4:**
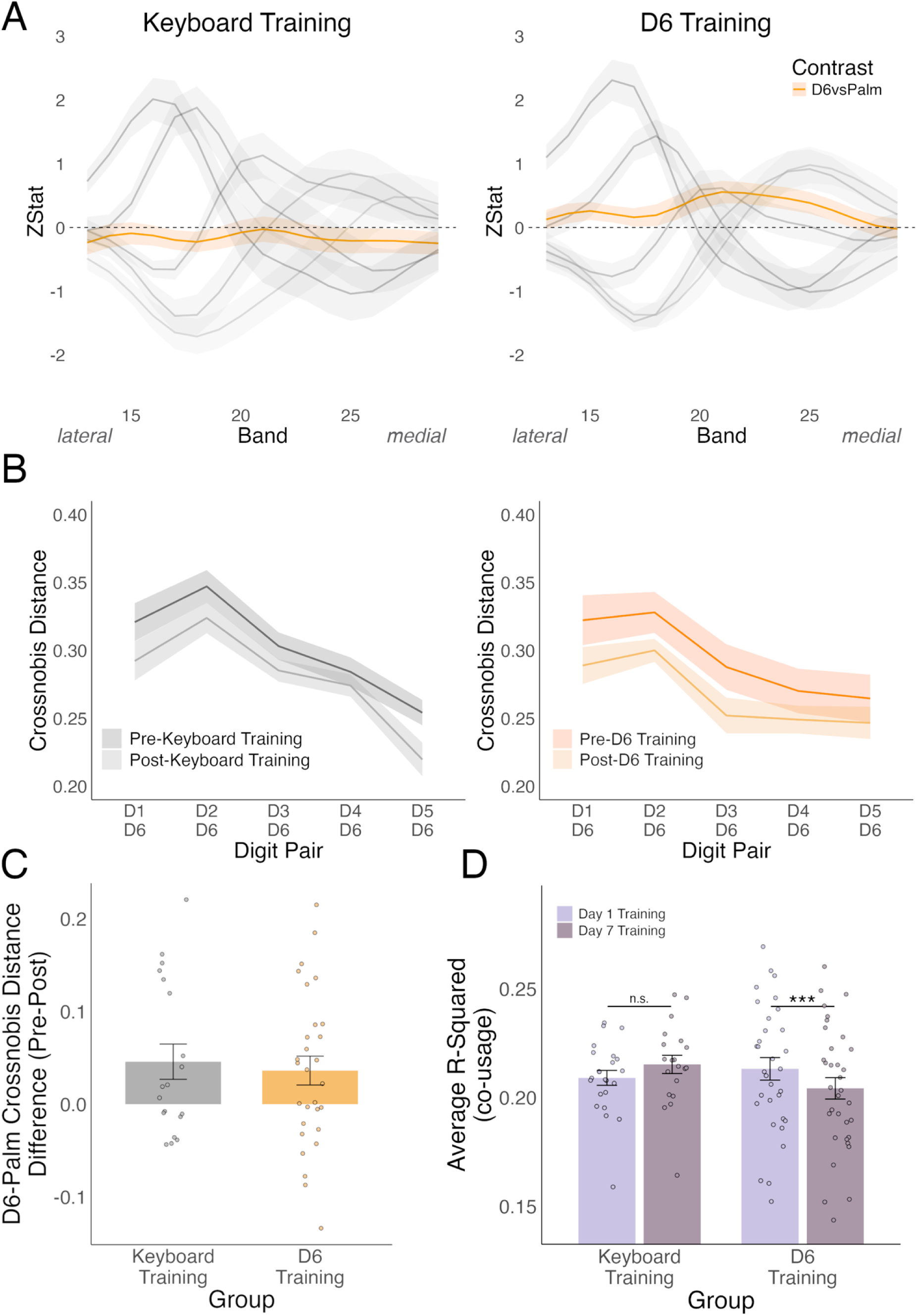
Refinement of the D6 sensory representation following D6 training. (A) Peak of activity for D6 within the S1 hand representation (versus palm). Following D6 training (right), we see a large and more distinct peak for D6, compared to the Keyboard control group (left). (B) We see reduced distances between D6 and the biological fingers for the D6 training group, relative to the Keyboard group, demonstrating that D6 is becoming more similar to the biological hand (C) The crossnobis distance between D6 and the palm does not change based on training experience (D) Co-usage measure of the biological fingers when wearing D6 and performing a series of free-choice tasks on the first and last day of training. We see the D6 group show increased finger individuation following training, not seen for the control group.

To quantify any refinement of D6’s neural representation induced by training, we assessed its similarity to (1) the palm and (2) the biological digits, using cross-nobis distance, across timepoints (pre/post training) and groups (D6/keyboard training). Relative to the palm, the D6 representation was stable (*F*(1,44)=1.032,*p*=.315; Figure 4C). However, the D6 representation became more associated with the biological digits, specifically following D6 training (group x session interaction: *F*(1,396)=8.08,*p*=.005; Figure 4B). Notably, this was found across all biological fingers, mirroring the behavioural results of similar gains across the whole hand. Put together, these results indicate that while the representation of D6 to the palm is context invariable, the rich interactions between D6 and the biological digits induced during training facilitated greater integration of D6 into the hand representation.

### D6-hand coordination training impacts hand kinematics

To better understand how D6 training affects this sensory integration, we compared finger kinematics during D6 task performance at the beginning and end of training. We measured biological finger co-use using a markerless tracking setup, during a series of free-choice tasks that required complex coordination patterns (see *46* for motor results; Figure 3D). Following longitudinal D6 training, we found decreased co-usage (i.e. increased individuation) of the biological fingers, which was not seen in the control group (group x session interaction: *F*(1,1950.73)=4.777,*p*=.029; Figure 4D). This suggests that prolonged D6 use induces changes in biological finger coordination patterns.

To assess whether these changes extend beyond direct D6 use, we also recorded finger kinematics while participants performed the same tasks using only their biological hand (D6 off). Following training, the D6 study group maintained the same co-use patterns whether or not they were using D6, whilst the control group showed a significant decline in finger co-usage (more free individuation) (group x D6 on/D6 off: *F*(1,1885.74)=13.33,*p*<.001; Figure 3E). This indicates that D6 training impacts hand kinematics beyond just the motor demands of D6 use; further implying D6 use may not only trigger changes to the D6 representation, but also of the biological fingers themselves (see *45*).

### Altered finger coordination impacts sensory hand representation

Given this impact on the biological hand, we also explored training-related changes in inter-finger representational relationships in S1 (disregarding D6), using RSA (see Figure S1 for univariate results). Both groups exhibited a significant reduction in inter-finger dissimilarity, reflecting an altered hand representation following sensorimotor skill learning (Figure 5), as previously reported for both D6 (*45*) and piano training (*53*,*54*). However, the piano control group showed a greater training effect, with a larger reduction in distances (group X session: *F*(1,836)=9.798,*p*=.002). These changes in neural representation compliment the kinematic effects, which demonstrated greater individuation in the biological hand-only free manipulation task (Figure 3D-E).

**Figure 5:**
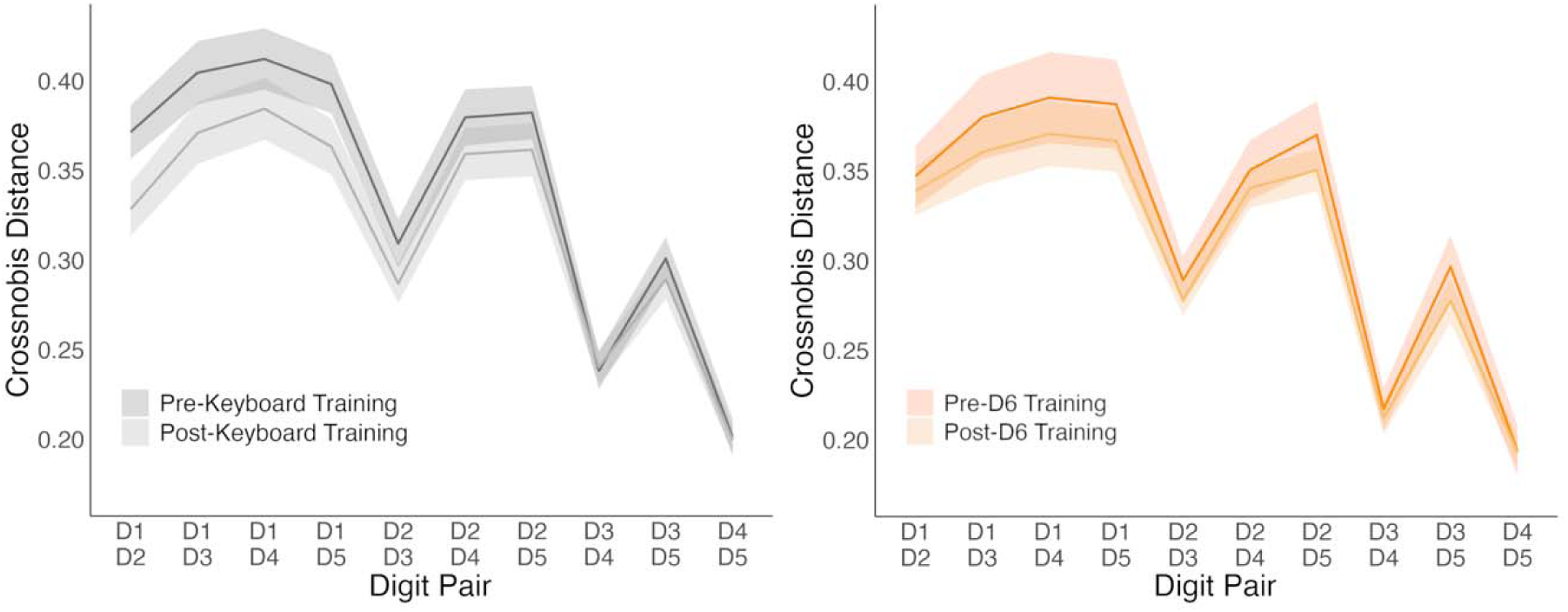
Effects of training on biological hand representation. Crossnobis distance for each group, before and after their respective training regimes (Keyboard left, D6 right). All participants demonstrated reduced inter-finger information content following their altered finger-coordination training, but we see a greater training effect for the Keyboard group.

### Neural integration of D6 is reflected by somatosensory embodiment following training

Finally, we examined whether D6 perceptual and neural integration was reflected in perceived (phenomenological) sense of embodiment. To assess this, participants completed a questionnaire in both the pre- and post-training sessions, covering multiple embodiment dimensions: Somatosensation, Body Ownership, Body Image, and Agency (latter discussed in *46*).

Before training, participants responded neutrally, and even negatively to embodiment statements across domains. Although participants could extract and integrate somatosensory cues from D6 (as reported in Figures 1C-E), this did not translate into subjective experience of somatosensory embodiment. Participants responded neutrally in the somatosensation category, with ratings not significantly different from zero (*t*(50)=1.527,*p*=.133,BF_10_=0.452). Responses to Body Ownership and Body Image were negative (significantly below zero, p=.018 and p<.001, respectively). Following D6 training, participants reported greater embodiment across categories compared to controls (group x session: *F*(1,226.85)=9.127,*p*=.003; Figure 6A). However, it was only in the Somatosensation category that D6 participants responded positively (significantly above zero; *t*(27)=3.840,*p*<.001; control group: *t*(18)=0.604,*p*=.553). For Body Ownership and Image, trained participants also responded neutrally (BF_10_=0.318 for both).

**Figure 6.**
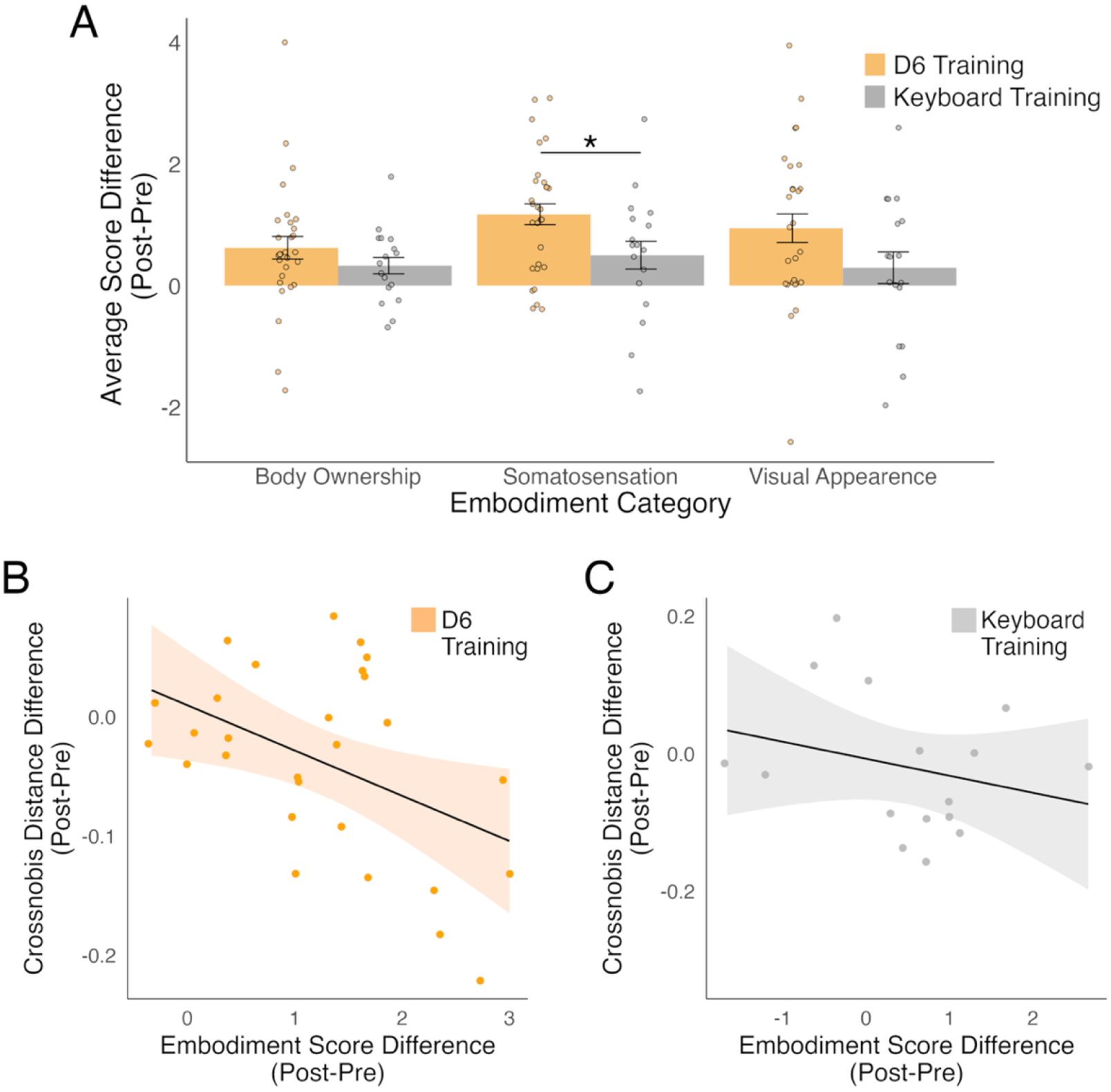
D6 training led to increased sensory embodiment. (A) Following training, D6 participants showed an increase in phenomenological embodiment compared to controls. Participants only experienced positive embodiment in the somatosensory category (consisting of questions such as “it seems like I can distinguish between different textures with the robotic finger” and “it seems like I can feel the touch of an object in the location with the robotic finger is touching the object”). (B) Significant negative correlation between decreases in cross-nobis distance (increased similarity) and increases in somatosensory embodiment scores (increased embodiment) D6 training, (C) Non-significant correlation between changes in cross-nobis distance and somatosensory embodiment for the Keyboard training control group. * denotes p < .05.

To establish if changes to experiential embodiment related to neural embodiment of D6, we correlated the change (post-pre-training) in cross-nobis distance between the D6-D1 pair, and in somatosensory embodiment ratings. The D6 training group showed a significant negative correlation (*r*(26)=-.451,*p*=.016; Figure 6B), not seen for the controls (*r*(14)=-.266,*p*=.320; Figure 6C), where decreased distance (increased similarity) between D6 and D1 corresponded to increased feelings of sensory embodiment (although note the two correlations were not significantly different from each other: *Z*=0.63,*p*=0.53).

Together, these findings support that extensive training with D6 facilitates somatosensory integration across all levels of the sensorimotor hierarchy-experiential, perceptual, kinematic, and neural.

## Discussion

Our findings provide a holistic characterisation of how the human sensorimotor system can integrate an artificial limb into its somatosensory body representation, offering insight into the plasticity mechanisms that underly how we perceive and interact with the world around us.

We have demonstrated that our somatosensory system can extract detailed information from the natural sensory cues arising from the physical interaction between an artificial limb (the Third Thumb (D6)) and integrate it with our body representation. Our body representation can quickly adapt to D6 across the sensorimotor hierarchy, suggesting we do not even need experience with an artificial limb to see meaningful integration with the body. However, this neural integration at the early stages of usage is not reflected in subjective experience; novice users had not developed a perceived sense of somatosensory embodied connection. Following training however, we observe multiple experience-dependent changes to hand-D6 integration. Not only do people successfully learn to use D6, but this experience also modulates adaptation of the body across both kinematic and neural matrices. These quantitative experience-driven changes in sensory integration appears to reflect a categorical shift in the subjective experience of the users, who now report feelings of somatosensory embodiment towards D6. These embodiment gains were asssociated with increased similarity between the way the biological hand and D6 are being represented in S1.

The fact that both the body and D6 representations are changing following extensive experience to use D6 calls for caution while considering the mechanism underlying this integration of D6 with the body. It remains an open question if this is a manifestation of sensory embodiment (*45*,*55*,*56*), or if this is simply reflecting the signatures of well-established adaptive plasticity mechanisms (*57*,*58*). When training to incorporate D6 into the motor repertoire, the body is undergoing two simultaneous changes – altered hand use, while also gaining a new somatosensory input (via D6). To disassociate the respective roles of these two changes, we can gain insight from our control group, who also trained to alter their hand use, but did not experience an additional somatosensory input. Indeed, while control participants show greater changes to the biological hand representation following training, the D6 training group showed increased changes to the integration of D6 into the hand representation. These results, which were also mirrored in the kinematics results, illustrates the unique role of sensorimotor experience in shaping the representation of an artificial body part into our body representation.

Nevertheless, it is important to highlight that the representation of D6 is inherently rooted in the biological hand representation – S1 does not reallocate neural resources of the body to develop an emerging representation of D6 (*39*,*40*). Instead, it integrates the inputs from D6 into our body representation. This observation distinguishes our findings from previous studies examining representation and embodiment of hand-held tools (*44*,*59–61*), and artificial limbs for substitution (*43*,*62–65*). Indeed, unlike tools, D6 is used *with* the hand, rather than *by the hand*. Therefore, the simultaneous changes to the representations of D6 and the biological digits result in a novel form of integration into the body representation, which sidesteps the issue of the limited neural resources that are available in S1 (*39*).

While our findings highlight the brain’s capacity for novel sensory integration, important questions remain regarding the purpose and longevity of these adaptations. A limit to our interpretation is that we do not know if the neural changes to the represenation of D6 observed in S1 are stable, and whether they serve a functional role. We previously demonstarted rapid learning with D6 (successful task performance within the first minutes of use (*66*)). Combined with the impressive performance of sensory tasks using D6 by naïve and novice participants shown here, it is possible that functional sensorimotor integration does not require an adapted D6 representation relative to the hand. Conversely, it is possible that further training could have evolved other differences to the S1 body and D6 representation. As augmentation technology becomes more embedded in everyday life, researchers must carefully consider its long-term implications for our bodies.

To conclude, we demonstrate the emergence of a distinct representation of an artificial body part within our body representation, and that this emerging representation is further integrated with the hand representation following sensorimotor experience. Understanding this malleability of our sensory body representation is crucial as we look to the future of technology, with successful sensory integration of a device being possible by the simple means of wearing it. By demonstrating the power of our somatosensory system to mediate and integrate such information, we have shown that you cannot change the body without changing the brain. This bidirectional relationship needs to be better harnessed when considering future technologies that interface with the human body and brain.

## Materials and Methods

### The Third Thumb

The Third Thumb (Dani Clode Design, London, UK) (Digit 6; D6) is a robotic extra digit that attaches to the ulnar side of the right hand (Figure 1A) and is operated through pressure sensors strapped underneath the big toes. D6 has two degrees of freedom that allow a corresponding proportional control: applying pressure to left big toe sensor causes an adduction/abduction movement, whilst pressure on the right one causes a flexion/extension movement (see further details in *45* and *67*).

### Feedback systems

#### Skin Stretch system

The skin stretch feedback system conveyed pressure information from D6’s tip to the user via a device worn on the inner wrist. The system comprised of three components: (1) a flexible pneumatic deformation sensor, integrated into D6’s tip, (2) an external control unit, and (3) a linear skin stretch device worn on the wrist (Figure 1C; Figure S2A).

A pneumatic deformation sensor was chosen for its compatibility with D6’s soft, flexible design. A silicone air chamber which deforms upon contact was integrated into D6’s tip, generating pressure changes measurable by the air pressure sensor. This tip was optimised using silicone moulding for improved sensitivity and durability. A custom circuit in the control unit processed the pressure signal, filtering noise before driving the skin stretch device. This device was positioned on the wrist for optimal sensitivity and used a high-torque servomotor to apply precise linear skin stretch, proportional to the pressure input. To enhance comfort and grip, the skin-contacting surface was lined with ethylene-vinyl acetate foam.

The system underwent multiple validation tests, where the pneumatic deformation sensor demonstrated 92% accuracy in classifying materials of varying deformability in a model hand test. The skin stretch device achieved 89% accuracy in a human material deformability 2AFC discrimination task involving three materials (full details available in supplementary materials).

#### Vibrotactile system

The vibrotactile feedback system conveyed information to the user about any displacement of the D6 tip caused by interactions with the external environment (predominantly object interactions), via high-frequency vibrations to the user’s ring finger. The system comprised of three components: (1) an inertial measurement unit (IMU) attached to the nail area of D6 for sensing, (2) a control unit mounted on the user’s wrist for processing signals, and (3) a vibrotactile actuator positioned on the back of the user’s ring finger to provide feedback (Figure 1D; Figure S2B).

The custom IMU mounted to D6’s ‘nail’ detected accelerations and vibrations caused by contact events at D6’s tip. The sensor was housed in a lightweight, 3D-printed protective casing. A custom electronic board processed the IMU data and generated control signals for the vibrotactile actuator. Signal noise was mitigated using a fourth-order Chebyshev Type I bandpass filter (120–230 Hz) which removed unwanted artifacts caused by free hand motion, and vibrations caused by D6’s servo and cable driven actuation system. The processed signal was mapped to a pulse-width modulation output, controlling the actuator’s frequency and amplitude. A linear voice coil actuator provided vibrotactile feedback proportional to the IMU output, encased in a custom 3D-printed housing. Elastic fabric within the housing ensured consistent vibrations by returning the actuator to its resting position after each pulse. The actuator was placed on the back of the ring finger, chosen for its high sensitivity to vibrations and minimal interference with hand function.

The vibrotactile system was validated using texture discrimination performance. Participants achieved 80% accuracy in a 2AFC task, demonstrating the delivery of interpretable and differentiable tactile information (see supplementary materials for full validation task details).

### Participants

In total, 102 participants took part in the study, see supplementary table S1 for breakdown of participants per sub-study. The study was approved by the Cambridge Psychology Research Ethics Committee (PREC: 2022.068) and the UCL Research Ethics Committee (project ID number 12921/001). Participants were recruited via online advertisements on the MRC Cognition and Brain Sciences Unit and University of Cambridge SONA participant pools.

### Material Discrimination Tasks

Using a within-subjects design, the same set of participants performed the material deformation and texture coarseness discrimination tasks across the three feedback systems, using a between-participant counter-balanced order. Due to the artificial feedback design requirements, we restricted the skin stretch system to the deformation task and the vibrotactile system to the coarseness task, whereas the wearable D6 system was used for both tasks.

#### General Task Design

Both material discrimination tasks were 2AFC. For the artificial feedback tasks, feedback was collected with a physically remote sensorised D6 and delivered via a worn actuator. For the natural feedback task, D6 was worn on the user’s hand during the tasks, and feedback provided via the intrinsic wearable interface. For the biological hand only task, the left biological thumb was used (left chosen due to task setup constraints).

In each trial, participants received a set reference stimulus first, followed by a comparison stimulus. In the deformation task, participants needed to determine if two deformable foam pieces were the same or different, and in the coarseness task, participants compared between two texture samples, based on the feedback cues available (natural or artificial feedback, in different blocks). Participants responded via a two-key keypad in their left hand.

Participants performed the task with their eyes closed, and with white noise playing through over-ear headphones. Volume was adjusted individually to mask the sound of a knock on the table. In addition to the white noise, participants were given a series of beeps to alert them prior to trial onset (t=-1.5sec), trial start time (t=0) and prompt to respond (t=5/3.5 for deformation/coarseness, respectively).

Participants were first familiarised with each task. They were asked to feel the various materials with their biological hand, and were then walked through an example trial to ensure task demands were clear. Participants were then given 10 pre-set practice trials with verbal reinforcement for correct performance. Each main experimental block included 50 trials with no verbal reinforcement.

In each trial, the comparison stimulus (the second stimulus delivered) was determined online using the adaptive psi method (*67*) implemented using the MATLAB Palamedes toolbox (*68*). The method estimates parameters of interest based on Bayes theorem, that are continuously updated after each trial response (*69*). We set broad prior estimates for both parameters of interest - slope and threshold (50%) based on pilot data. Psychometrics parameters were preset as follows: Deformation discrimination: threshold (−20 – 20; 0.01 steps); slope: −2 – 2; 0.1 steps); Coarseness discrimination: threshold (−10 – 10; 0.01 steps); slope: −2 – 2; 0.1 steps)

Offline analysis involved fitting the responses for each task/system block to a psychometric logit curve and calculating the Akaike information criterion (AIC) as a measure of goodness of fit. We compared this to the AIC of the intercept-only (no predictor) model and excluded any datasets showing a difference <2 (*70*). In total 8.05% of the data was excluded due to fit issues (see supplementary table S1 for breakdown per block). We next extracted the slope of the fitted curve for each block/participant, as a measure of sensitivity while discriminating between the different samples, with a larger value indicating better discrimination ability. We also replicated the analysis with the values of the intercept, informing us about the bias people have in responding (see supplementary materials).

#### Material Deformation Task

Each material piece was cut to the approximate size of 8.5 cm x 6 cm x 4 cm. Eight out of the nine materials were foams varying in deformability, the ninth was a solid block. Deformability was initially quantified using the average signal peak when a deformation sensor embedded into the D6 tip sampled each foam piece three times. The most deformable material had a measure of 0.7 and used as reference, whilst the firmest material had a measure of 29 (full range: 0.7, 2.2, 5.2, 8.9, 10.8, 11.8, 15.1, 19.1, 29). The smaller the value, the less force was received in response when the deformation sensor sampled the material. Deformability order was later corroborated by quantification in cm through an in-house system that delievered 850g of weight to each foam (amount of deformation: 2.6cm, 2.4cm, 1.5cm, 0.8cm, 0.75cm, 0.65cm, 0.45cm, 0.2cm, 0)

Task setup is presented in Figures 1C and S4. The hand (wearable device) or model hand (skin stretch) was secured underneath a table that allowed full flexing of D6 to sample the materials, while minimising contact with the hand. D6 was triggered to automatically move to sample the material (never controlled by the participant), hold for one seconds, and then release. In the biological thumb-only control task, the participant timed the sampling themselves, practicing beforehand.

#### Texture Coarseness Task

All the texture pieces were 79 mm long and 35 mm wide and made from silicone, ensuring compatibility with the D6 tip. Each texture had a series of pyramid shaped peaks, creating finer or coarser ridges. The nine textures were quantified by the distance between peaks. The finest texture had a 5mm distance between peaks (used as reference), whilst the coarsest had a 15mm distance between peaks (total range: 5mm, 6mm, 7mm, 8mm, 9mm, 10mm, 11mm, 13mm, and 15mm).

Task setup can be seen in Figures 1D and S5. The hand (wearable device) or model hand (vibrotactile) was secured on a mount off the table to allow sampling of the texture pieces by D6, while minimizing contact with the hand. D6 was always passive in this task (never connected to power so could not move), it was simply worn on the hand/hand model whilst the texture pieces were applied to the tip, using a custom rail controlled by the experimenter.

### Temporal Order Judgement

#### Hardware

To deliver the vibrotactile stimulations, encapsulated DC vibration motors (3 VDC) were attached to the glabrous skin on the fingertips of the biological thumb (D1), index finger (D2), little finger (D5) and the silicone pad of D6 (Figure 1D; Figure S6). Each stimulation lasted 15 ms, with a frequency of 235Hz and vibration amplitude of 7G.

#### Task Design

The task comprised of four digit-pair blocks, delivered in a randomised order – D6&D1, D6&D2, D6&D5 and D5&D1. Each block had 40 trials; in each trial participants received vibrotactile stimulation to one digit and then another in quick succession (interstimulus interval (ISI): −400ms to 400ms, possible steps of 10ms between). Participants verbally reported which digit they felt was stimulated first (the digit to the left of the digit-pair (respond “0”) or the digit to the right (respond “1”) and were reminded of the response mapping at the beginning of each block.

Before the task began, participants completed six practice trials for each digit-pair at pre-determined ISIs. Here participants had full vision and audition and received verbal reinforcement for correct performance.

In the experimental run, participants had their eyes closed and listened to white noise through over-ear headphones; The white noise was initially set to 75% of the maximum volume, with adjustments made to ensure comfort. They received no verbal reinforcement.

The interstimulus interval between stimulations was determined using the adaptive psi method (described above). The threshold estimate was set to −0.1 to 0.1; steps of 0.01, with a slope estimate of −1.5 to 1.5; steps of 0.1.

We replicated the curve fitting and goodness of fit checks described above for each digit-pair block, leading to removal of 1% of the data. We calculated the JND as a measure of discrimination ability by finding the difference between the 75% point and 25% point on the psychometric curve and dividing by two. Smaller values imply better temporal discrimination ability. We also extracted the point of subjective equality as a measure of bias by finding the 50% point of the psychometric curve, to explore spatial biases (supplementary materials).

### Training

The full protocol for both training groups is available in the general protocol at *osf.io/c76xd*.

#### Third Thumb (D6) Training

Participants in our experimental group underwent seven days of D6 training, designed to allow acquisition of different motor skills with D6, particularly focused on D6-biological hand collaboration. The first and last day of training was supervised in-person, whilst the middle five days were completed remotely and semi-supervised (approximately two hours of training per day). Full details available in *46*.

On the first and last day of training participants performed a D6-biological digit coordination task. Per one-minute block, participants had to make tip-to-tip movements between D6 and a randomised digit (cues: ‘thumb’, ‘middle’, ‘ring’, ‘little’ – four blocks). Participants were given the audio cue ‘go’ to begin a trial, and ‘stop’ to end the trial when the experimenter determined the coordinated movement was successful.

#### Keyboard Training

Our active control group underwent a similar training regime (two days in-person, five remote, with approximately two hours of training per day). Participants trained to play the piano keyboard over seven days of training, guided by the app ‘Yousician’ (*71*), using only their right hand and a right foot pedal to maintain the notes.

Participants completed one set of songs in the app at the beginning of every session, and a different set at the end of every session. In between, participants completed different training tasks working on technique, major scales, minor scales and blues scales. Each training task was repeated 10 times each, replicating the D6 training schedule (both schedules available at *osf.io/c76xd*). Yousician calculated a score for each task based on note-accuracy and timing information. Participants were given a different chord to practice in between each session.

### Functional Magnetic Resonance Imaging (fMRI)

#### Acquisition

Details of all MRI images acquired are available at *osf.io/c76xd*.

MRI images were acquired using a 3T Prisma Fit MRI scanner (Siemens, Munich, Germany) using a 32-channel head coil. For functional image acquisition, we used a multiband T2* - weighted pulse sequence with a between-slice acceleration factor of 4 and no in-slice acceleration. A voxel size of 2mm isotropic was used, and a repetition time (TR) of 1500 ms. Acquisition sequence parameters consisted of an echo time (TE) of 35 ms, flip angle of 70° with 72 transversal slices per volume. For our T1-weighted image acquisition sequence to acquire our structural image we used a voxel size of 1mm isotropic and TR of 2250 ms, with sequence parameters consisting of a 3ms TE, flip angle of 9°.

#### Task and tactile stimulation

Silicon vibrotactors were used to deliver vibrotactile stimulations via a pneumatic system to the glabrous skin of the five biological fingertips, to the silicon tip of D6, and to the side of the hand where D6 is worn (the ‘Palm’) (Figure 2A). We used an in-house system inspired by the design of *72* (validated in *73*). The digit vibrotactors were held in place with elasticated sports tape, whilst the Palm vibrotactor was fitted under the D6 hand piece.

During the scan, each body part was stimulated individually in a pseudo-randomised order. Each tactile stimulation event was delivered over a nine second block – a one second preparation period for system pressure stabilisation, 7.5 seconds of stimulation, then a 0.5 second rest of no stimulation. To avoid peripheral or central adaptation (*73–76*), each stimulation varied frequency every 400 milliseconds, following a pattern of 5 Hz, 15 Hz, and 30 Hz, with 100 millisecond gaps between frequencies; this pattern occurred five times per block. Interweaved within the stimulation blocks were six, nine second ‘rest’ blocks with no stimulation, to allow for BOLD signal relaxation. Rest blocks could be on their own or grouped in two consecutive blocks, but between rest block groups there was always at least two stimulation blocks. In addition, two 16 second rest blocks were at the beginning and end of each run.

Participants completed three runs of the task, each 472 seconds long (323 volumes). There was no active task, but to ensure participant engagement they were told (a) to ensure the name of the digit on the screen matched the digit being stimulated and (b) to inform the experimenter if any vibrations in the run felt different to the others. When available, an eye tracker was used for a subset of participants to ensure attention was maintained throughout the task.

#### Pre-processing and first-level analysis

All MRI data pre-processing and analyses were carried out using FMRIB Software Library ((*77*); FSL, version 6.0.5), as well as in-house scripts written in MATLAB (version R2020b), Python (3.13.0), and R (4.2.2). For multivariate analyses, we used the MATLAB RSA Toolbox (*78*) in addition to within-lab toolbox extensions (*79*). The decoding analyses were carried out using python library scikit-learn (v.1.5.2).

To ensure that the functional scans were well aligned for each participant, we calculated a midspace between the six runs (three pre-training runs, three post-training runs). This represents the average space where the images are minimally reoriented. Each scan was aligned to this midspace using FLIRT (*80*,*81*), with structural image alignment optimised using Boundary-Based Registration.

Functional data was primarily pre-processed using FSL-FEAT. We first performed motion correction using FMRIB’s Linear Image Registration Tool (MCFLIRT (*81*), followed by brain extraction using the Brain Extraction Tool (BET (*82*)). We then used temporal high-pass filtering, with a cutoff of 100 seconds, and lastly spatial smoothing with a 3 mm full width at half maximum (FWHM) Gaussian kernel. Field maps were also used for field unwarping.

We used a voxel-based general linear model (GLM) implemented in FSL-FEAT to identify activity patterns related to each tactile stimulation condition. For each run, seven regressors were created, one for each body part condition (D1-D6, Palm). Regressors were convoluted with a double-gamma function to account for the delayed BOLD response. Additional first derivative regressors were included to capture temporal variations in the BOLD signal, along with six motion parameters to estimate head movements. Additionally, we used motion scrubbing, adding regressors of no interest to exclude any volumes with excessive motion (framewise displacement larger than 0.9 (*83*)). In the model, 14 contrasts were set up; each body part vs rest, each biological digit vs all the other biological digits, D6 vs Palm, and the average of the biological digits vs rest. For each participant, the contrast images obtained for each of the three runs per session were then averaged voxel-wise using a fixed-effects model in FSL. This image was normalised to the standard Montreal Neurological Institute (MNI) space and used for the *Line Analysis*. For the Line Analysis, we focused on the selectivity in each condition by contrasting the biological digits and D6 relative to their respective controls - the average of the other digits, and the palm of the hand.

#### Region of Interest (ROI) Definition

We focused all analyses in S1, specifically Broadmann Area 3b (BA3b). For each participant, their structural T1 image was used to estimate the cortical surface, reconstructing the pial and white-grey matter surfaces using FreeSurfer’s recon-all (Freesurfer, 7.4.0 (*84*)).

To initially explore selectivity across the BA3b strip, we took the standard flat map (FS_LR 32K) of the Human Connectome Project (HCP) and performed a 15° clockwise rotation so that the central sulcus was perpendicular to the horizontal axis. We then created a mask of BA3b as defined by the Glasser Atlas (*85*), and constructed 50 equally spaced lines from the top to bottom of the mask (anterior to posterior axis), creating 49 ROI ‘bands’. These were then transformed into volumetric ROIs. We constrained then visualisation to the estimated hand area, bands 13-29.

To localise the hand representation for further analyses, we took the Glasser BA3b ROI and a pre-defined ROI taking all surface S1 nodes on the standard flat map 2cm proximal/distal of the anatomical hand knob (*86*). We transformed this into volumetric standard MNI space and took the intersection of the two ROIs. We further refined the ROI by taking the top 100 most active voxels for each biological condition contrasted to rest (each biological digit vs rest, and palm vs rest) from the activity of an independent group (*N*=20) that completed the paradigm. We took the additive sum of these voxels as our final ROI. These were then mapped to the individual’s volumetric functional space (via the individual’s volumetric high-resolution anatomy). An important consideration is that this ROI may not precisely reflect BA3b and may contain relevant activity from neighboring S1 areas due to the nature of our data and processing (3T fMRI, smoothing FWHM 3mm) and the probabilistic nature of the atlas. As such, we take this as definitively S1, and indicatively BA3b, as such as refer to this as S1 throughout.

#### Representational Similarity Analysis (RSA)

To explore the informational content available in the hand representation, we used multivariate representational similarity analysis (RSA (*87*)). For each participant and each session, we extracted the beta weights estimates from the first-level analyses for each condition vs rest within our BA3b hand area ROI. These were prewhitened using the GLM residuals, and the cross-validated mahalanobis (crossnobis) distance (*88*) was calculated between the patterns for each of the conditions. We calculated the distances using each possible imaging run pair combination within a session (pre- and post-training) and then averaged the distances across all run pairs. This distance provides us with a measure of dissimilarity in the activity patterns elicited in each stimulation condition. The expected distance is zero if two patterns are not significantly different from each other. The crossnobis distance between D6 and D1 was taken to correlate with embodiment scores due to being the D6-biological digit pair most commonly used in collaboration (*46*) and the pair that would intuitively have the largest distance based on hand topography.

### Markerless Tracking

We extracted the hand use kinematics during a set of free-choice tasks that engaged complex digit coordination patterns (see task details in *46*) using a markerless tracking setup. We recorded performance using three Logitech Brio 4k Stream Webcams (*89*). During offline analyses, we tracked the 2D joint coordinates of the five biological digits using Google Mediapipe (*90*) for each of the three camera recordings. We then filtered and triangulated using Anipose to obtain the 3D joint angles (*91*). The intrinsic parameters of the cameras and the relative location were obtained during calibration using a ChArUco board (Size 21.0 x 29.7 cm; 10 x 7 squares, with 4 bit markers and a dictionary size of 50 markers).

For each participant, session and task, the joint angles were extracted and smoothed using third-order Savitzky-Golay filter with a window length of 151 samples. Angular velocities were then calculated from the first difference of the filtered joint angle data divided by the time step (*45*). We then z-normalized across the tasks within a session (*92*). Separately for the distal interphalangeal joint (DIP) and proximal interphalangeal (PIP) joint angles (using the interphalangeal joint (IP) for D1), we used linear regression to fit the angular velocity data of a given digit as a function of the angular velocity of each of the other digits individually. This produced a coefficient of determination (R^2^) for each digit pair, informing about the proportion of total variance of each digit’s angular velocity that could be explained by a linear reconstruction, based on the paired regression with each of the other digits. We took this as a measure of co-usage.

### Embodiment Questionnaire

Participants completed an Embodiment questionnaire at the end of their pre-training and post-training behavioural sessions to assess phenomenological experience with D6. The questionnaire contained 20 statements (adapted from *45*; based on *93*), which participants rated along a seven-point Likert scale, ranging from −3 (strongly disagree) to 3 (strongly agree).

Statements formed four categories – somatosensation, body ownership, body image, as well as agency (reported in *46*). There were additional four ‘catch’ statements, designed to confirm the participant was paying attention to their ratings. For each participant and session, questionnaire scores were averaged within each category. If a participant’s average rating in the catch category exceeded zero, their data for that session was excluded, resulting in the removal of one participant.

### General Analysis

All statistical analyses were performed in JASP (version 0.19.1). Throughout, parametric statistics were used when there were no violations of test assumptions (normality assessed by Shapiro-Wilk values and visual inspection, homoscedasticity assessed by Levene test values). If there were violations, when possible, the equivalent non-parametric test was used. All null results of interest were followed up using Bayesian statistics (Cauchy prior width r=0.707) to obtain a Bayesian factor (BF_10_) to allow interpretation based on the strength of support for the null hypothesis. We used a threshold of BF<1/3 as sufficient evidence in support of the null hypothesis, BF>3 as sufficient evidence in support of the alternative hypothesis, and 1/3<BF<3 as inconclusive evidence (*94*).

Wherever possible, we employed linear mixed models (LMM), with a random intercept for participant. We used the Satterthwaite testing method and Satterthwaite degrees of freedom approximation for all follow-up tests. Models accounted for testing session, group and digit-pair (fMRI multivariate analyses and kinematics) or category (embodiment questionnaire) as fixed effects. Our fMRI models additionally accounted for days between sessions using a continuous covariate, and our kinematics models accounted for joint (PIP or DIP).

For any simple comparisons where a LMM would not be appropriate, we used one-sample t-tests to assess if values were significantly different from zero (including the material discrimination tasks, fMRI multivariate analyses, and embodiment questionnaire) and paired samples t-tests and repeated measures ANOVAs for within-subjects comparisons (including material discrimination tasks and TOJ performance). For both, a Wilcoxon signed-rank test was used if assumptions were violated. For the between-groups comparison when comparing material discrimination task performance with our biological hand to the wearable D6 performance, a Mann-Whitney test was used due to assumption violations. A Pearson correlation was used to correlate embodiment and Crossnobis distance.

## Acknowledgements

We thank all of our participants for their time and effort. We thank Raffaele Tucciarelli for support with experimental setup and analysis; Ilana Nisky for helpful advice during development of the feedback systems; Silvestro Micera and Solaiman Shokur for helpful advice throughout the study; Yousician for providing us with a preminum version of their application for this research; Hristo Dimitrov for support with setting up the kinematics analysis; Mabel Ziman & Clara Gallay for data collection; and Roderick Spender, Klara Selén, Katarina Krajnovic, Hannah Browning, Yuval Amichay, Karan Salvi and Alexandra Williams for additional support with data collection. We thank Yon Visell and Calogero Oddo for their helpful feedback on the initial manuscript.

## Author Contributions

L.D and T.R.M led all conceptualisation and experimental design development. M.M and G.D supported experimental design for the longitudinal training participants. D.C designed and constructed the Third Thumb. E.d.S led design and construction of the artificial feedback systems. M.B, F.I, D.C. L.D and T.R.M supported artificial feedback system development. M.B and F.I were also available for consulation throughout. L.D contributed to data collection for all the datasets included in this manuscript. M.M, G.D, E.d.S, E.J and V.P also contributed to data collection on subsets of the data included in this manuscript. L.D and T.R.M led all data analysis. G.D and V.P. contributed to organisation and processing of the kinematics data. E.J. contributed to processing and analysis of the keyboard training data. M.M and V.P. contributed to organsiation and processing of the Third Thumb training data. L.D and T.R.M wrote the paper. L.D and D.C prepared the figures. M.M, G.D, E.d.S, V.P, E.J, M.B, F.I, and D.C. provided feedback on the paper. T.R.M provided funding for the paper.

## Data and materials availability

The data that support the findings of this study will be available from the Open Science Framework upon publication (osf.io/c76xd). For the purpose of open access, the authors have applied a Creative Commons Attribution (CC BY) licence to any Author Accepted Manuscript version arising from this submission.

## Supplementary Materials

### Supplementary Results

#### No differences in response biases between artificial and natural feedback in material discrimination tasks

In addition to the slope, we also extracted each participant’s threshold (50% point on the fitted psychometric curve) for each block/participant to examine response biases. For the Material Deformation task, both feedback systems had values significantly above zero (*W*≥209,*p*<.001), implying a bias towards responding the samples were different. Although, this is likely due to the higher number of different trials (*t*≥4.671,*p*<.002). Participants did not show bias differences between the artificial and natural feedback conditions (*W*(19)=148,*p*=.114,BF_10_=.911).

For the Texture Coarseness task, we again found a bias towards responding that the samples were different (*W*≥171,*p*<.001), likely due to the higher number of different trials (*t*≥8.48,*p*<.001). However, participants had a larger bias for reporting the textures were different in the artificial condition (*W*(15)=24,*p*=.021), and there was no difference in the number of different trials (*t*(15)=.232,*p*=.819,BF_10_=.262). Considering the reduced sensitivity to differences in the artificial condition (Figure 1D), this bias suggests participants compensated for increased uncertainty by adopting a response strategy favoring ‘different’ judgments.

#### No differences in spatial biases when integrating D6 sensory information

For the TOJ task, we calculated the point of subjective equality (PSE) to explore spatial biases in responses (a positive or negative PSE indicates a bias towards responding that the digit to the right or left (respectively) was stimulated first).

The biological digit-pair did not significantly differ from zero (*W*(48)=720,*p*=.290,BF_10_=.187), implying no biases in responses with natural sensing. D6 biases were then comparable, not differing from zero (*W*≥506, *p*≥.295, BF_10_≤.226) nor from the biological digits (*W*(48)=524.00, *p*=.385, BF_10_=.251). We also did not find differences between the three D6 digit-pairs (*W*(2)=.005,*p*=.775,BF_10_=.124), suggesting distance did not impact how the sensory information from D6 was integrated with the digits.

Following training, we saw no changes in biases across all four digit-pairs (session: *F*(1,328.36)=0.302,*p*=.583; higher-order interactions also non-significant (*p*≥.613)). Considering there was no difference from the biological digits even in novice users, this implies that from early exposure D6’s perceptual representation accurately represents the true state of the world, and this does not shift with experience.

#### No changes to perceptual integration following D6 training

The lack of change in perceptual integration following training is further evidenced by the lack of change in discriminability ability, indexed using the just noticeable difference (JND). Again there was no significant change in JND following training, across all digit-pairs (session: *F*(1,328.23)=1.201,*p*=.274; higher-order interactions non-significant (*p*≥.492)). Before training, participants already performed comparably with the D6 and only the biological digits, so this lack of change implies participants had already reached ceiling.

### Supplementary Methods

#### Skin Stretch System and Feedback Validation

To validate the functional capabilities of the pneumatic deformation sensor at the tip of D6, a material deformation discrimination task was employed. This task utilised four different materials: three types of foam with varying deformation values and fully solid material (PE500).

The sensor was fully integrated into an actuating D6 that was attached to a model hand and power supply. A foam was placed in the model hand, and D6 was actuated to flex (80% flexion) and sample the foam 50 times for each of the materials. This data was then input into a decision tree classifier (Classification Learner App; MATLAB R2022a) which showed 92% accuracy in distinguishing between the four materials (Figure S3A).

The Skin Stretch system was validated through pilot testing (*n*=5). They engaged in a preliminary version of the material deformation task (2AFC). Data previously gathered from three materials (the least dense foam, a median density foam, and a solid material) during the sensor validation phase were utilised to generate pre-determined inputs for the device. For each material, average maximum and minimum values were calculated and were used to define a specific angular range (percentage of skin stretch) induced by the device. For each stimulation the amount of skin stretch for a material varied randomly within this predefined range to mitigate learning effects during the task.

Participants wore the skin stretch device on their inner wrist, they closed their eyes and wore noise-cancelling headphones emitting white noise to eliminate visual and auditory feedback. In the task, they received two sequential skin stretch stimulations, as a pair (one after another) with an interstimulus interval of 2s. They were then asked to determine whether the stimulations were the same or different. Each participant completed 60 trials, comprising an equal number of ‘same’ and ‘different’ pairs (30 same, 30 different). The results showed that using the skin stretch device, the participants were able to distinguish between the three different materials with an overall accuracy of 89% (Figure S3B).

#### Vibrotactile System and Feedback Validation

To validate the actuator output from the vibrotactile system, we used an external IMU that allowed recording of vibrotactile output. D6, integrated with the vibrotactile system, was secured to a flat surface via a clamp that. A manually operated sliding rail and carriage system delivered textures to D6’s tip with uniform pressure and consistent timing via a beep-cue. A sample consisted of a texture piece being slid across the tip of D6 in one direction at a constant speed, the resulting vibrations were collected by the external IMU. 50 samples were collected for three different texture sizes (5mm, 9mm and 15mm). This version of D6 had a single ridge tip.

The data was classified using a coarse decision tree classifier (Classification Learner App; MATLAB R2022a). The device reached a high 70% accuracy rate (Figure S3C), despite using data derived through a secondary IMU, which is subject to a considerable amount of noise.

Another texture coarseness discrimination task was set up to identify the ideal position to provide the vibrotactile feedback on the user’s body. Three different texture pieces (5mm, 9mm, and 15mm) were used in 2AFC paradigm, where the participants were provided with two texture stimulations in a row (a pair) with an inter-stimulus interval of 2 seconds. Participants had to verbally respond if they felt the same or different. D6, with the integrated vibrotactile system, was set up on the clamp and texture slider setup, and participants wore the vibrotactile actuator, where they received the feedback following the texture delivery. All participants closed their eyes and wore noise-cancelling headphones playing white noise to block out visual and auditory feedback.

Participants (*n*=5) tested this paradigm with the actuator placed on three different body position: fingertip of the index finger (baseline), back of the ring finger and the elbow joint bone. 60 trials were delivered per body position, with an equal number of same and different texture pairs.

The fingertip of the index finger would serve as a benchmark for the highest possible accuracy, due to it having the highest density of mechanoreceptors in human skin (*95*,*96*). The index finger was not considered as a final site to deliver the feedback, as it would hinder use of the hand. The back of the ring finger was selected as a possible location due to its sensitive skin (attributed to a higher density of mechanoreceptors compared to the arm), proximity to bone, enhancing vibration sensitivity, and its location not hindering hand functionality. The elbow, characterized by its thin skin layer and direct bone contact, offers excellent vibration sensitivity. Its placement on the arm ensures hand functionality remains unaffected.

Participants could perform with the feedback delivered to any of the location, but the back of the ring finger emerged as the optimal location, achieving an accuracy of 80% (matching index finger performance; Figure S3D), and we therefore selected it as the final position for the vibrotactile feedback.This decision led to a redesign of the final iteration of the actuator, specifically tailored to fit the back of the ring finger, ensuring optimal performance and user comfort.

#### Participant demographics and exclusions

**Table S1:** Participant demographics split by task and training subset, details for all additional exclusions.

| Task | *N* | Age | Gender | Additional Exclusions |
| --- | --- | --- | --- | --- |
| Material Discrimination | *N*=22 | *M*=24.96<br>(*SD*=3.86) | 13 Female,<br>9 Male | *N*=1 material deformation task with *artificial feedback* due to technical issues<br>*N*=1 material deformation task with *artificial feedback* due to poor fit<br>*N*=4 texture coarseness task with *artificial feedback* due to poor fit<br>*N*=2 texture coarseness with *natural feedback* due to poor fit |
| TOJ | *N*=50 | *M*=24.7<br>(*SD*=4.17) | 35 Female,<br>18 Male | *N*=2 at pre-training from D6D5 digit-pair due to poor fit<br>*N*=1 at pre-training from D6D2 digit-pair due to poor fit<br>*N*=1 at pre-training from D5D1 digit-pair due to poor fit |
| fMRI | see TOJ | see TOJ | see TOJ | *N*=1 ended scan early due to participant illness<br>*N*=1 due to consistently negative cross-nobis distances |
| Kinematics | see TOJ | see TOJ | see TOJ | *N*=1 due to missing calibration<br>*N*=4 due to failed calibrations |
| Embodiment | See TOJ | See TOJ | See TOJ | *N*=1 due to missing data, additional<br>*N*=2 missing pre-training, *N*=1 missing post-training<br>*N*=1 at pre-training due to positive scores on catch trials |
| D6 training | *N*=30 | *M*=24.7<br>(*SD*=3.91) | 19 Female,<br>11 Male |  |
| Piano Keyboard training | *N*=20 | *M*=24.7<br>(*SD*=4.57) | 16 Female,<br>7 Male |  |

**Figure S1:**
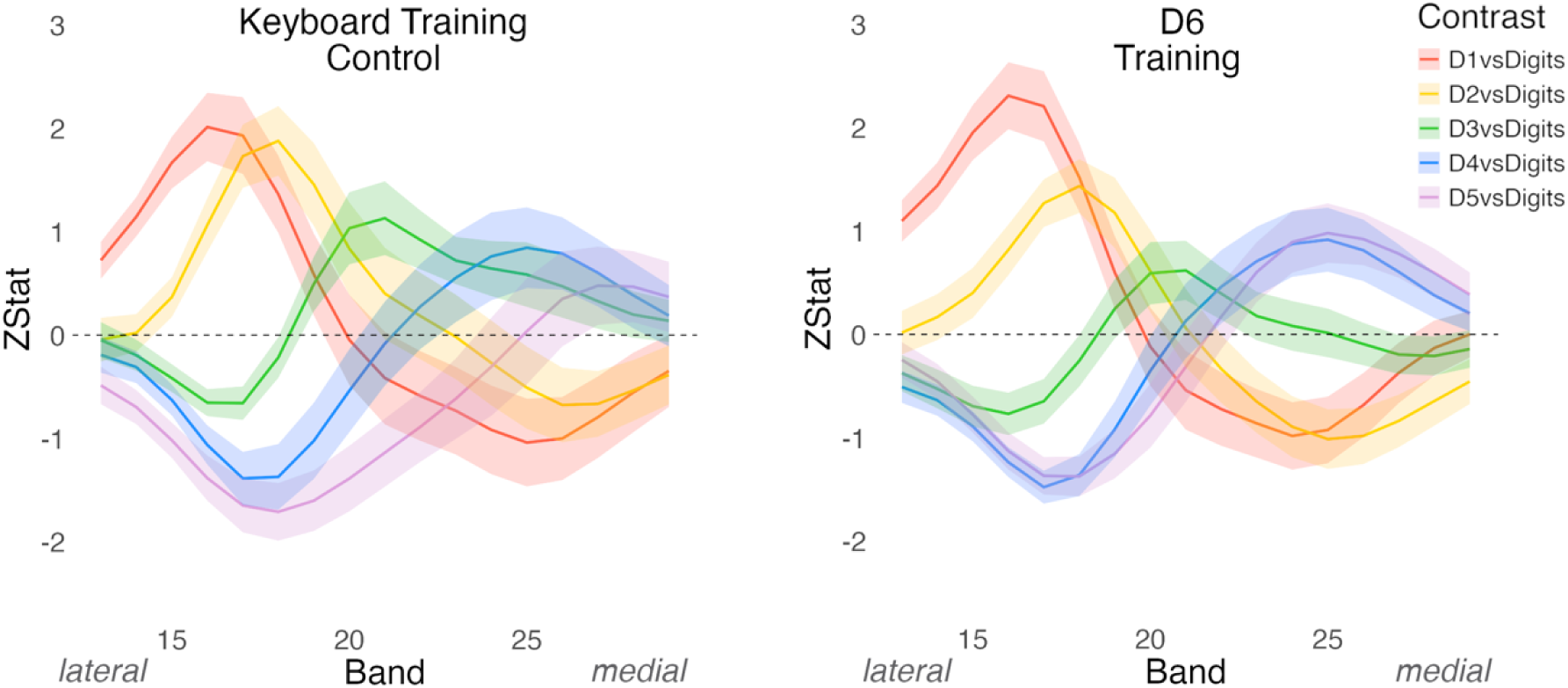
Univariate activity of the biological digits with the hand representation post-training. (A) Peaks of activity for each of the biological digits across the S1 hand representation following Keyboard training (left) or D6 training (right), from lateral (left) to medial (right).

**Figure S2:**
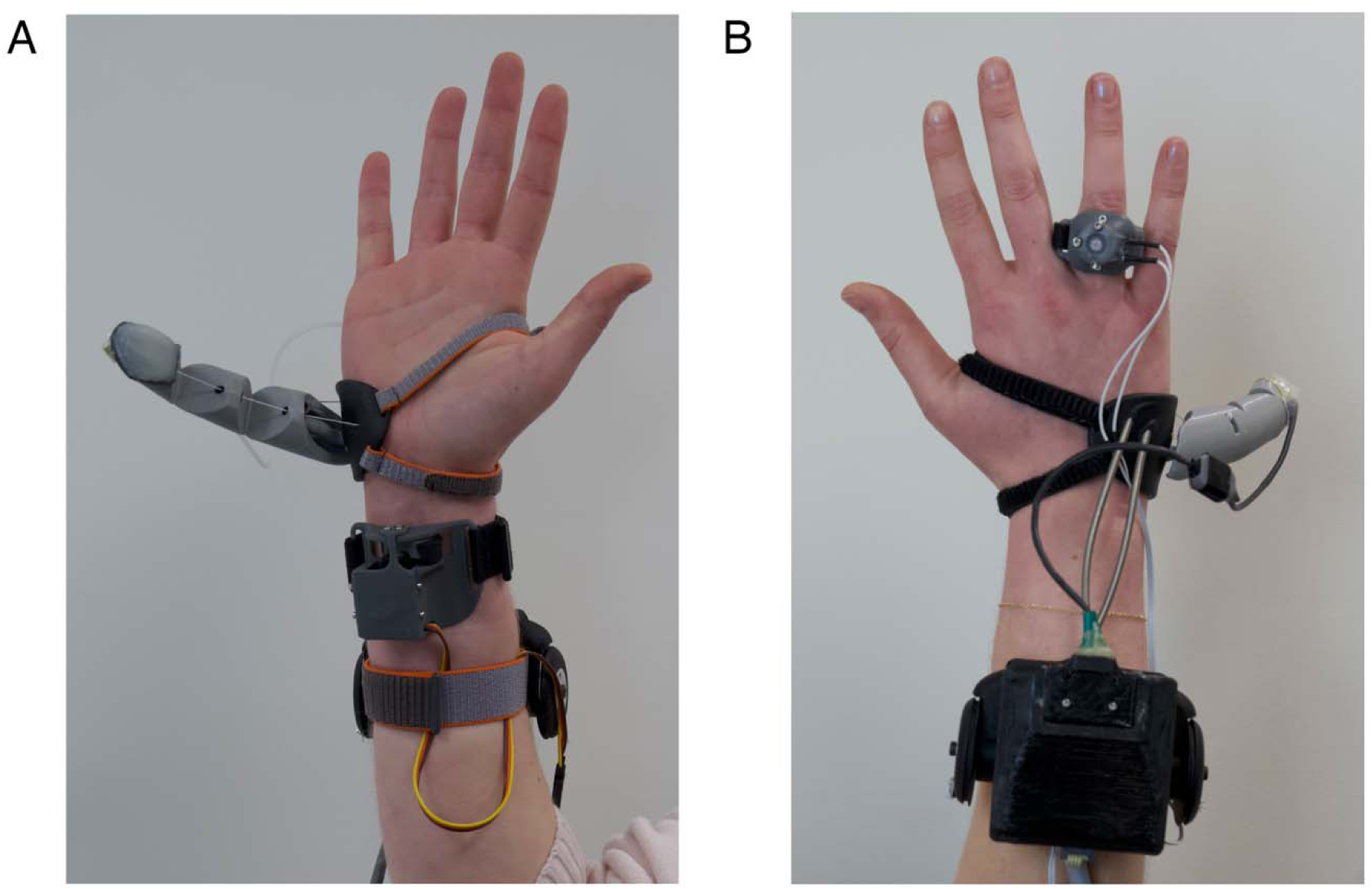
Artificial feedback systems integrated with D6. (A) Skin Stretch feedback D6 system. A flexible pneumatic deformation sensor is integrated into D6’S tip, a linear skin stretch actuator is worn on the inner wrist (B) Vibrotactile feedback D6 system. An inertial measurement unit is attached to the nail area of D6 for sensing, a vibrotactile actuator is positioned on the back of the user’s ring finger

**Figure S3:**
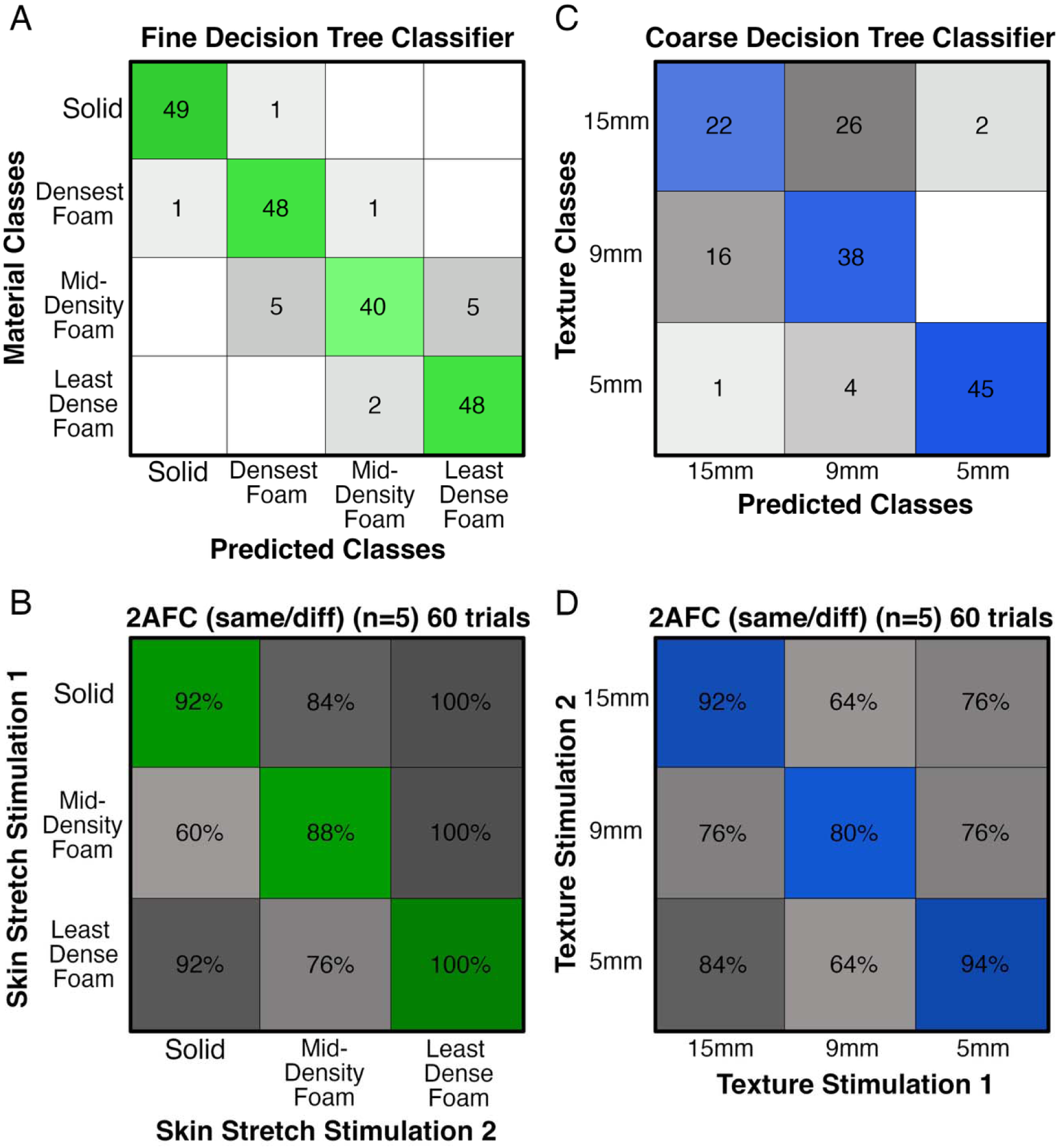
Artificial feedback validation tests. (A) Material classifications from a fine decision tree classifier, classifying the skin strech feedback signals from foam samples, achieving 92% accuracy (B) Participants’ performance when discriminating the skin stretch signals, achieving 89% accuracy (C) Texture classifications from a coarse decision tree classifier, classifying the vibrotactile feedback signals (via an IMU) from the texture samples, achieving 70% accuracy (D) Participants’ performance in a texture coarseness discrimination task using the vibrotactile feedback signals, achieving 80% accuracy when delivered to the back of the ring finger.

**Figure S4:**
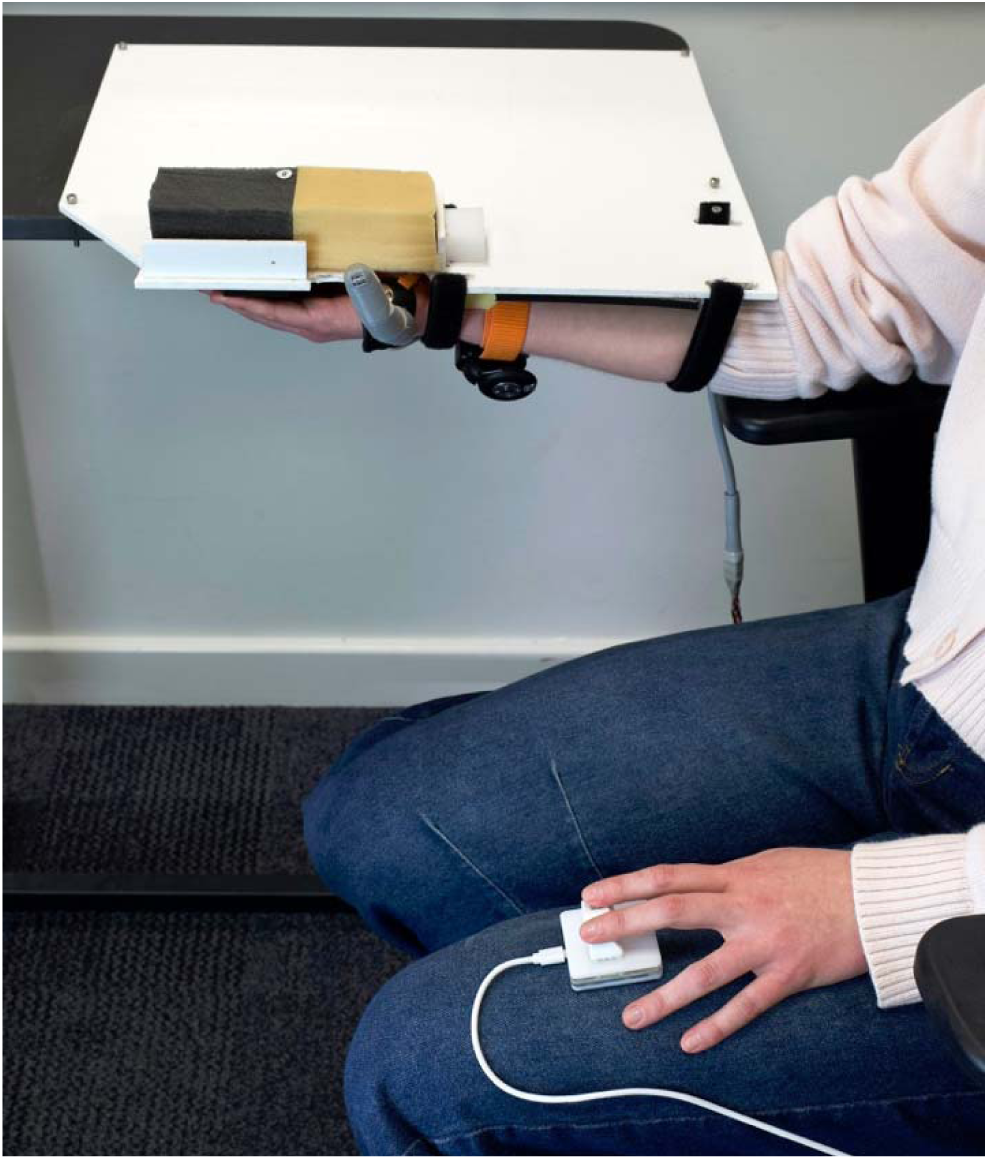
Material deformation task setup. Full setup for the Material deformation task. Triggers were sent to automatically move D6 to sample different foams, and the participant (with closed eyes and headphones playing white noise) made perceptual judgements based on the feedback they received, responding via a keypad

**Figure S5:**
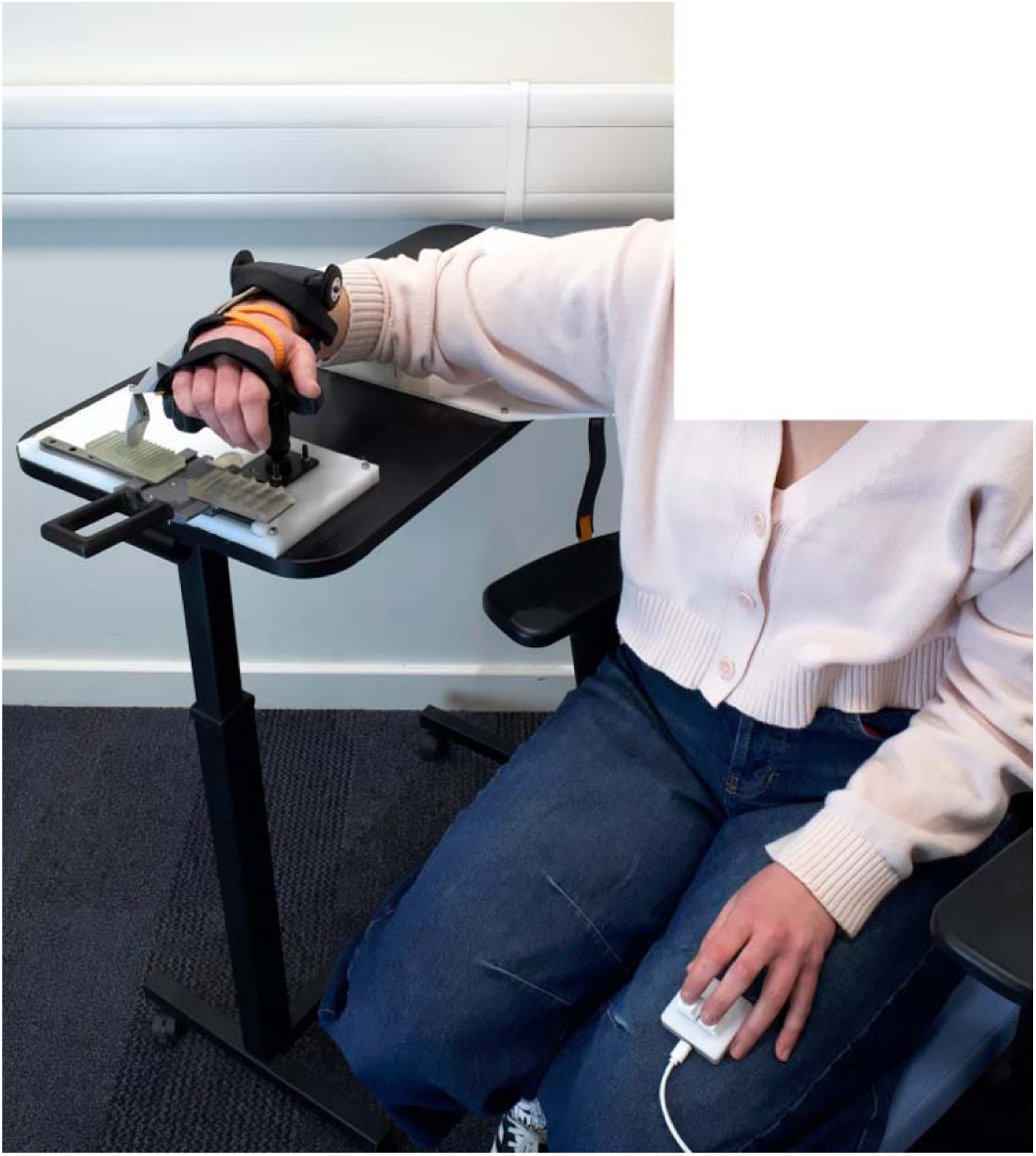
Texture coarseness task setup. Full setup for the Texture coarseness task. Textures pieces were moved along a rail by the experimenter for D6 to sample, and the participant (with closed eyes and headphones playing white noise) made perceptual judgements based on the feedback they received, responding via a keypad

**Figure S6:**
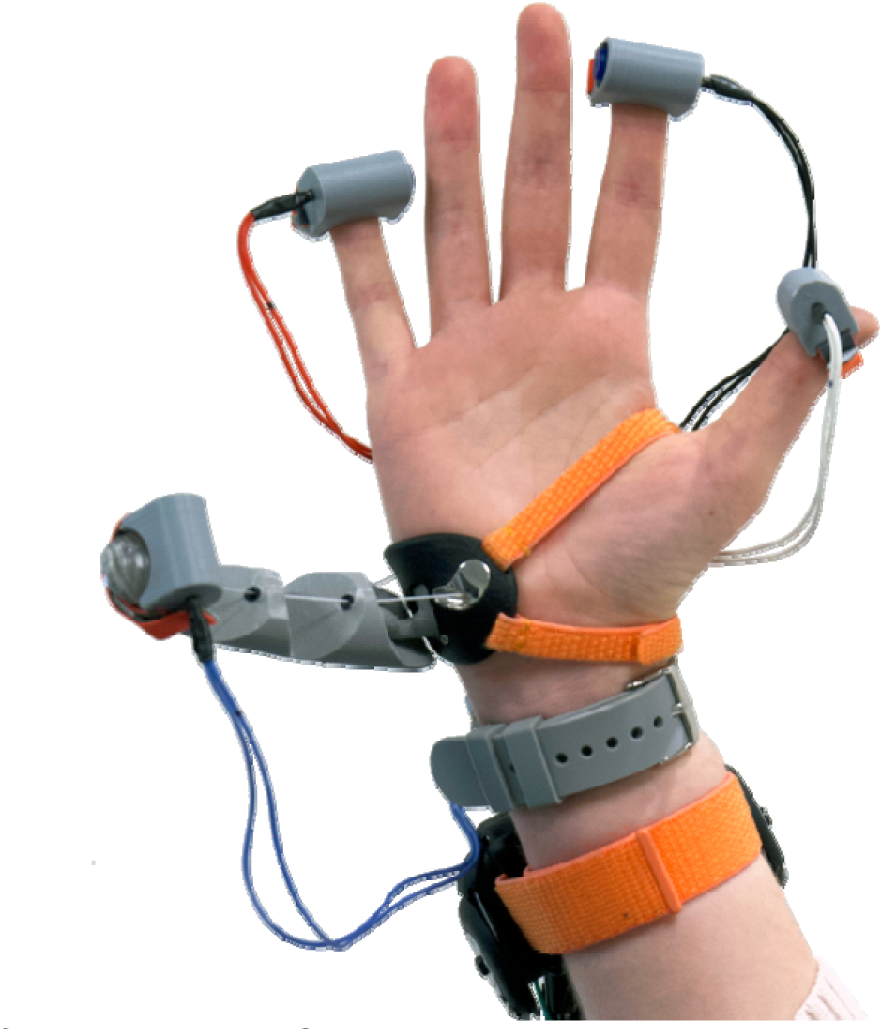
Full setup for Temporal Order Judgement task. Motors attached to the biological thumb (D1), index finger (D2), little finger (D5) and Third Thumb (D6).

